# A Note about the Individualized TMS Focality

**DOI:** 10.1101/2020.02.10.941062

**Authors:** Sergey N. Makarov, William A. Wartman, Mohammad Daneshzand, Aapo Nummenmaa

## Abstract

A particular yet computationally successful solution of an inverse transcranial magnetic stimulation (TMS) problem is reported. The goal has been focusing the normal unsigned electric field at the inner cortical surface and its vicinity (the D wave activation site) given a unique high-resolution gyral pattern of a subject and a precise coil model.

For 16 subjects and 32 arbitrary target points, the solution decreases the mean deviation of the maximum-field domain from the target by a factor of 2 on average. The reduction in the maximum-field area is expected to quadruple. The average final deviation is 6 mm.

Rotation about the coil axis is the most significantly altered parameter, and the coil moves 10 mm on average during optimization. The maximum electric field magnitude decreases by 16% on average. Stability of the solution is enforced. The relative average de-focalization is below 1.2 when position/orientation accuracies are within 3 mm/6 degrees, respectively. The solution for the maximum normal field may also maximize the total field and its gradient for neighboring cortical layers III-V (I wave activation).

## 1. Introduction

Noninvasive, noncontact transcranial magnetic stimulation (TMS) [1]–[3] uses electromagnetic induction to generate electric fields internal to the brain remotely via a coil placed next to the subject’s head. It enables painless and safe suprathreshold stimulation. One common site of TMS activation is the primary motor cortex. The TMS response of the primary motor cortex – a muscle twitch used for motor mapping, particularly in preoperative settings [4], for assessing spinal cord injury [5]–[7], and recently for brain computer interfaces [8] – is measured with implanted cortical electrodes [9]–[11] or via electromyography (EMG) [12]–[14]. Another common activation site is the dorsolateral prefrontal cortex of the left hemisphere, which is pathophysiologically linked to depression [1]–[3].

The TMS activation mechanisms are complex. The TMS response includes the D (or direct) TMS wave and multiple I (or indirect) waves [9]–[11]. The earliest D wave originates from direct stimulation of pyramidal axons of the fast-conducting pyramidal tract neurons in the subcortical white matter or in the axon initial segment [10],[11]. The I waves, which occur 1.2–2.0 ms after the D wave, originate from indirect, trans-synaptic activation of neurons [10]–[14]. The I waves are more common at low suprathreshold field intensities. The D wave is recorded at higher suprathreshold field intensities and/or for specific (lateral-medial) coil orientations [10]–[14].

Compared to intracortical electric microstimulation (cf. [15],[16]) and, more recently, to transcranial focused ultrasound [17], TMS is thought to suffer from poorer targeting and focality. In particular, the apparent TMS motor map may extend due to remote hotspot activation [18]. TMS focality has previously been investigated from the viewpoint of a better TMS coil design, better coil penetration depth, and better field concentration in Refs. [19]–[23]. None of these studies investigated individualized TMS focality resulting from the unique gyral pattern shape.

Very interesting recent development is to align the TMS coil to the individual central sulcus or elsewhere for better, muscle-specific representation of the motor hand area [18],[24]–[26]. Since the TMS activation domain is determined by the electric field gradient distribution in the gyral pattern of the neocortex, we suggest that accurate, high-resolution field modeling might enable us to significantly improve the TMS focality for an individual subject once the detailed brain model is available. This is the task of the present study. This task is formalized as an inverse problem: find the best stable coil position/orientation for targeting a specific point in a specific cortical domain. The inverse TMS problem in a basic form has been considered in Ref. [23].

To make the inverse problem solvable, the target cortical domain and the field type must be defined. In this study, we restrict ourselves to one domain – the white-gray matter interface and its vicinity, as well as to the corresponding normal field. This domain is the source of D wave activation in either straight [28], or bent [27],[28],[29] pyramidal axons, or in the initial segment of axons [11] of fast-conducting neurons from layer V. Furthermore, the maximum interfacial normal electric field in this domain well correlates with the maximum total electric field.

The core of the inverse-problem solution is a direct solver capable of resolving rapidly varying electric fields in the cortex with the maximum precision and, simultaneously, capable of solving many (thousands) of possible field distributions in a reasonable amount of time. We apply a new boundary element method formulation in terms of induced surface charge density at the interfaces naturally coupled with the fast multipole method [30],[31] or BEM-FMM [32],[33]. The method possesses high numerical accuracy, which was recently shown to exceed that of the comparable finite element method or FEM [34]. Once the solution for the induced surface charge density is obtained, the normal electric field just inside or outside any brain surface is found precisely and without postprocessing. Most important, the method provides numerically unlimited field resolution close to and across the cortical surfaces. The BEM-FMM resolution is not limited by the FEM volumetric mesh size and may reach a micron scale if desired.

## 2. Materials and Methods

### 2.1. Targeted TMS response – D wave

The earliest D wave measured with implanted electrodes [9]–[11] or surface EMG electrodes [12]–[14] originates from direct stimulation of pyramidal axons of the pyramidal-tract neurons in the subcortical white matter or in the axon initial segment [9],[11]. The D wave is recorded

i. at higher suprathreshold field intensities [10],[11],[13],[14], e.g. above the active motor threshold [9]–[11] and up to 150% of the active motor threshold [14] and/or;
ii. for specific coil orientations (e.g. lateral-to-medial field direction) [10],[11],[14].

In this study, the earliest D wave of the TMS response is targeted. Its anticipated activation site – the white-gray matter interface and its immediate vicinity – is well-localized in space and could be accurately segmented and tracked using available high-resolution (0.5-1.0 mm isotropic) T1/T2 MRI datasets and modern automated segmentation software [35],[36].

### 2.2. Targeted cortical domain – white-gray matter interface and its vicinity

The targeted domain schematically shown in Fig. 1 by a pink color is the white–gray matter interface and its immediate vicinity. This domain is the most likely activation site of the D wave following three possible axon activation mechanisms discussed in the literature: (i) a rapid change of the normal electric field across the interface [28] and the resulting high value of the activating function [38],[39]; (ii) strong axonal bending in the vicinity of the interface [27],[28],[29] and the resulting high value of the activating function [38],[39]; and (iii) a strong field in the initial segment of pyramidal axons [11] originating from the fast-conducting neurons of layer V.

**Fig. 1.**
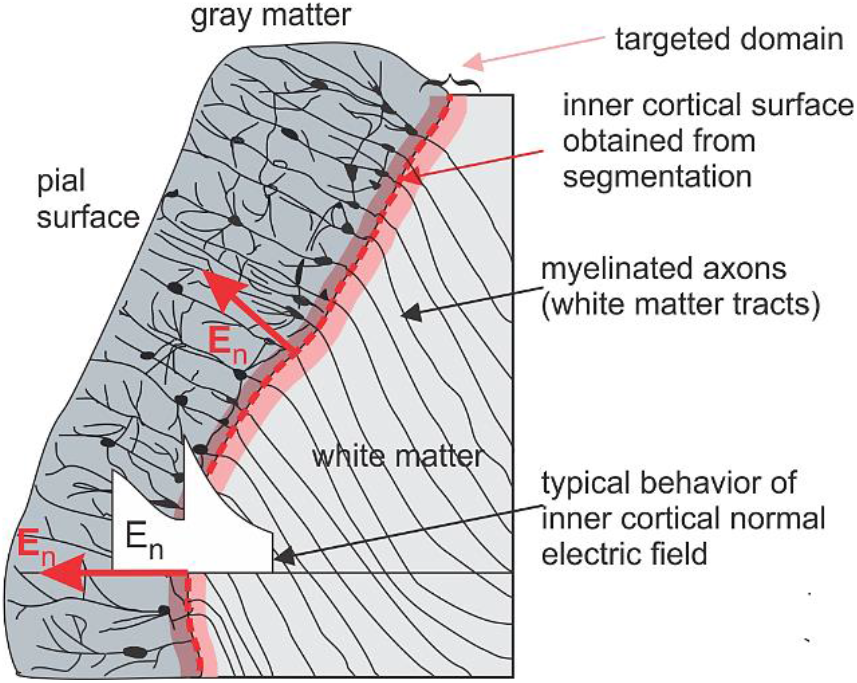
An illustration highlighting the targeted domain: the white-gray matter interface obtained from image segmentation and its immediate vicinity marked by a pink color. Typical behavior of the normal electric field, *E*_*n*_, across this interface is shown by a white graph. *E*_*n*_ undergoes a rapid change when passing the interface, its value just inside (white matter) is significantly higher than the value just outside (gray matter).

### 2.3. Targeted field – normal field at the white-gray matter interface

Due to a high conductivity contrast between gray matter (GM) and white matter (WM), with the conductivity ratio exceeding two at common TMS pulse widths [37], a significant surface charge density, *ρ*(***r***), will reside at the interface [28]. We distinguish the normal electric field just inside the interface, *E*_*n,in*_(***r***); the normal electric field just outside the interface, *E*_*n,out*_(***r***); and the normal field discontinuity for any conducting interface, Δ*E*_*n*_. It can be show using the method of surface integral equation that, for any observation point ***r*** ∈ *S* at the interface, for any primary TMS field, and for any head model,

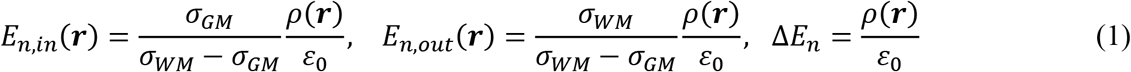

where *ε*_0_ is the permittivity of vacuum and *σ*_*GM*_ > *σ*_*WM*_ are the corresponding conductivities. Typical behavior of the normal electric field, *E*_*n*_, across this interface is shown in Fig. 1 by a white graph. The field just inside (white matter) is significantly higher than the field just outside (gray matter). All three quantities in Eq. (1) are proportional to each other and are proportional to the induced charge density. Therefore, we will simply refer to all of them as the normal field in the following text. Note that the maximum normal electric field at this interface correlates with the maximum total electric field as shown in the Discussion section.

In fact, the infinitely thin white-gray matter interface in Fig. 1 is an abstraction; it indeed has a finite thickness. For example, the myelination that is important for the electrical insulation of the axons starts to increase already in cortical layer IV and below in the rat neocortex [40]. The finite yet small thickness value should not alter the solution given above significantly.

### 2.4. Targeted cortical areas – motor hand area and dorsolateral prefrontal cortex of 16 Connectome subjects

The two targeted cortical areas are the motor hand area, namely the hand knob [41] of the right hemisphere, and the dorsolateral prefrontal cortex of the left hemisphere, which is pathophysiologically linked to depression [1],[42]–[44]. MRI data for sixteen Human Connectome Project (HCP) healthy subjects [46] with an initial isotropic voxel resolution of 0.7 mm have been converted to surface models using the SimNIBS v2.1 pipeline [35]; each model includes seven brain compartments (skin, skull, cerebrospinal fluid (CSF), gray matter (GM), white matter (WM), ventricles, and cerebellum) [47]. Each model has been checked against the original NIfTI images, and mesh manifoldness has been strictly enforced and confirmed using ANSYS HFSS mesh checker. The average cortical surface mesh edge length is 1.5 mm, the cortical nodal density is 0.55 nodes per mm^2^, and the total average number of facets is 0.9 M.

### 2.5. TMS forward problem solution – boundary element fast multipole method (BEM-FMM)

The TMS forward problem is the computation of the electric field in the head (including the cortex) given the coil geometry, position, and orientation. We use the boundary element method formulated in terms of induced surface charge density *ρ*(***r***) residing at the conductivity interfaces and naturally coupled with the fast multipole method [30],[31] or BEM-FMM, originally described in [32],[33]. The method possesses high numerical accuracy, which was recently shown to exceed that of the comparable finite element method of the first order [34]. Once the solution for the induced surface charge density is obtained, the normal electric field at any interface is found precisely and without postprocessing (cf. Eq.(1)).

Most important, the method supports unconstrained field resolution close to and across cortical surfaces, including both the outer cortical surface (the interface between gray matter and cerebrospinal fluid following terminology of [45]) and the inner cortical surface (the interface between white matter and gray matter). Since the solution is fully determined by the conductivity boundaries, the BEM-FMM field resolution within the cortex is not limited by the FEM (finite element method) volumetric mesh size and may reach a micron scale if desired.

### 2.6. Inverse problem formulation. Search space. Constraints. Cost function

The nonlinear inverse problem considered here is an attempt to produce a relative “hot spot” (or spots) of the normal electric field at the inner cortical surface localized near a target point by moving and rotating a TMS coil as shown in Fig. 2.

**Fig. 2.**
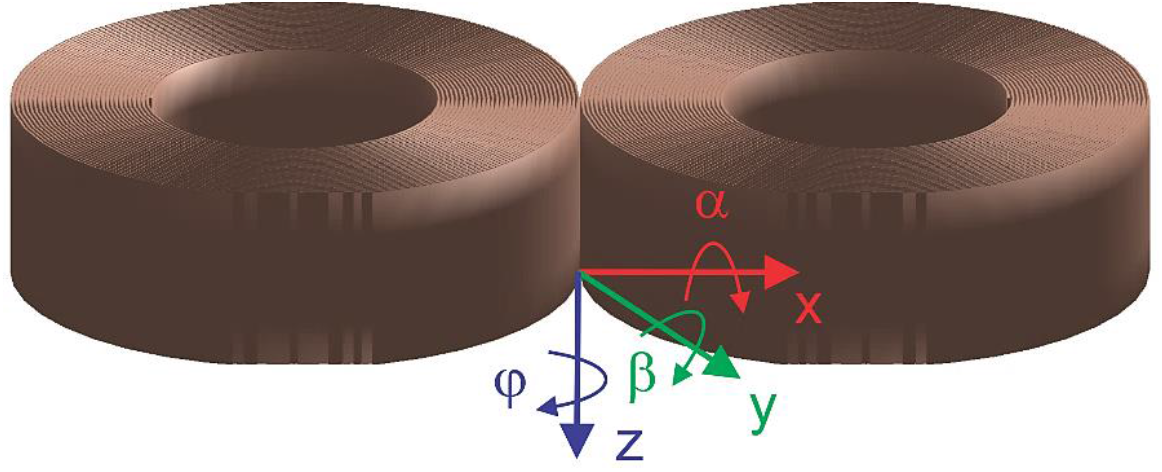
Three rotation angles of a TMS coil under study: angle *φ* about the *z*-axis (yaw); angle *α* about the *x*-axis (pitch); angle *β* about the *y*-axis (roll). The commercial coil model used in this study is MagVenture Cool B35 coil (approximated by 120,000 current dipoles via FMM [48]).

The commercial coil model used in this study is a modern compact figure-eight MagVenture Cool B35 coil. The coil current distribution is approximated by 120,000 elementary current dipoles via FMM [48].

Given anticipated clinical applications, the target point itself will be defined at the outer cortical surface (the gray matter shell). This is done to enable better visual navigation of the subject’s gyral pattern.

The search space is a six-dimensional space (ℜ^6^) that includes three Cartesian coordinates of the coil center and three coil rotation angles (pitch-roll-yaw) shown in Fig. 2. The only constraint imposed implies that the *nearest* distance from any part of the coil to the skin of the subject under study is no less than 10 mm.

The cost function is defined as the mean or average absolute deviation (ADD) of the field maxima from a target point ***T*** in 3D, that is

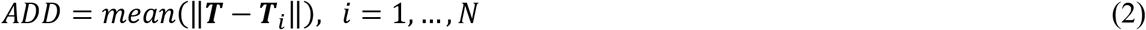

where ‖·‖ is the Euclidian norm, and ***T***_*i*_ are centroids of *N* white matter facets where the absolute normal-field value just inside the white matter shell is in the range of 80-100% of the maximum normal-field value just inside the same surface. Identical hot spots would also be observed for the normal component just outside the interface and for the normal field discontinuity, Δ*E*_*n*_, across the interface. The measure of Eq. (2) does not square the distance from the target, so it is less affected by a few extreme observations than the variance and standard deviation.

### 2.7. Initial guess

For each of 16 subjects, the spatial target point ***T*** was arbitrarily selected within the approximately identified hand knob area of the right precentral gyrus. Similarly, the target point, ***T***, was arbitrarily selected within the dorsolateral prefrontal cortex of the left hemisphere. The initial coil placement (the initial guess) follows three rules:

i. align the target point, ***T***, with the coil centerline;
ii. set the coil’s centerline perpendicular to the skin surface at the skin-centerline intersection, and position the coil bottom 10±0.25 mm from the skin surface along this centerline;
iii. Do not enforce lateral-to-medial or anterior-to-posterior field directions a priori. Instead, have the dominant coil field direction be approximately perpendicular to the gyral crown (and the associated sulcal walls) of the particular gyrus that contains the target point.

### 2.8. Search algorithm

The cost function has been minimized in the following manner. First, an exhaustive search algorithm is employed to solve the problem for 3^6^ = 729 neighbor coil positions/orientations in the ℜ^6^ search space around the initial guess (labeled with subscript 0). The 729 initial search points are given by all combinations of:

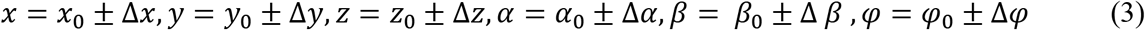

where we choose Δ*x* = Δ*y* = Δ*z* = 2 mm and Δ*φ* = Δ*α* = Δ*β* = 0.1 *rad* = 5.7°. The cost function is computed for every search point, and the search point with the absolute cost function minimum is chosen as the next guess in Eq. (3). Further, the process repeats itself for no more than 12 consecutive steps. When the convergence saturates (the current guess coincides with the previous guess), the Δ variations in Eq. (3) are automatically increased by a factor of 1.2. This search method is a combination of a local exhaustive search algorithm with a gradient-based correction. Its advantage is an ability to track the global minimum of the cost function.

## 3. Results

### 3.1. Solution of the inverse problem for the motor hand area of the right hemisphere

Fig. 3 shows a typical solution of the inverse problem for subject #110411 of the Connectome database. Fig. 3a corresponds to the initial guess while Fig. 3b is the solution obtained after 5 iterations. The target point ***T*** is indicated by a magenta ball. The spread of the intracortical normal electric field within 80-100% of the maximum field is indicated by small blue balls on the white matter surface. In the first case (Fig. 3a-right), the spread of the field involves not only the targeted motor hand area of the precentral gyrus but also the postcentral gyrus and further the superior parietal lobule. This situation is rather typical for conventional TMS. In the second case (Fig. 3b-right), the field is much more focal: the 80-100% peak field is well localized around both sulcal walls of the precentral gyrus and is close to the target point. The final cost function in Eq. (2) is equal to 7.0 mm.

**Fig. 3.**
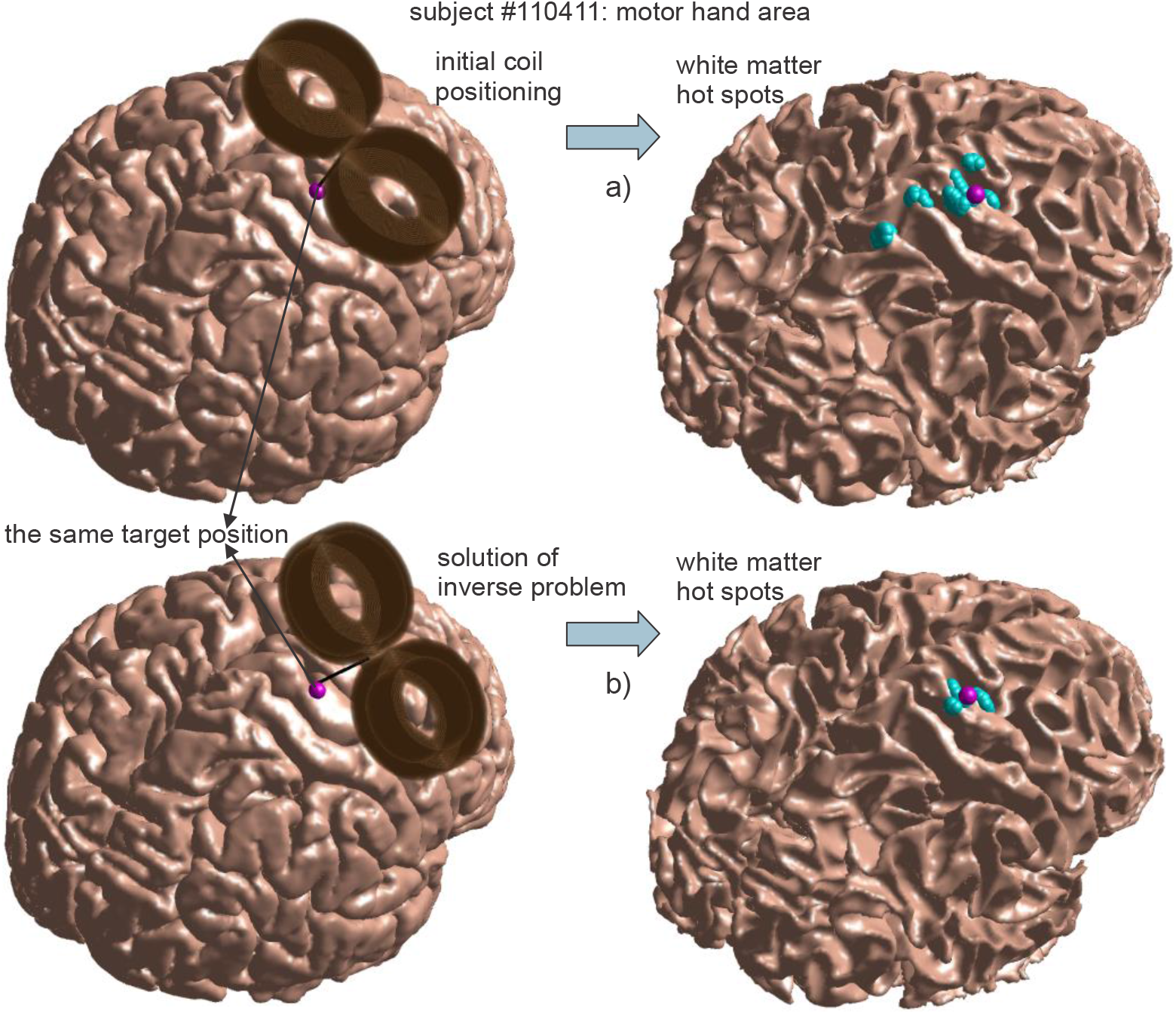
a) – Left: initial coil positioning 10 mm above the motor hand area of the right hemisphere for subject #110411 (GM surface is shown). The target position on the gray matter surface is marked by a magenta sphere. Right: white-matter “hot spots” – small blue balls drawn at the center of every WM facet where the absolute normal-field value is in the range of 80-100% of the maximum normal-field. b) – The same as a) but the coil position on the left is altered according to the inverse-problem solution.

Similar results have been obtained for the remaining 15 cases. Every solution was further checked for stability and corrected (if necessary) as described in the Discussion section.

### 3.2. Solution of the inverse problem for the left dorsolateral prefrontal cortex

Fig. 4 shows a typical inverse problem solution for the left dorsolateral prefrontal cortex. We again consider the same subject #110411 of the Connectome database. Fig. 4a corresponds to the initial guess, while Fig. 4b is the solution obtained after 8 iterations. The target point ***T*** is indicated by a magenta ball. The spread of the intracortical electric field within 80-100% of the maximum field is indicated by small blue balls on the white matter surface. In the first case (Fig. 4a-right), the spread of the field involves not only the targeted part of the superior frontal gyrus but also the middle frontal gyrus. This situation is again typical for conventional TMS. In the second case (Fig. 4b-right), the spread of the field is smaller: the 80-100% peak field is well localized around the superior frontal sulcus wall of the targeted gyrus and is very close to the target point. The minimum cost function in Eq. (2) is 4.9 mm. Similar results have been obtained for the remaining 15 cases. Every solution was further checked for stability and corrected (if necessary) as described in the Discussion section.

**Fig. 4.**
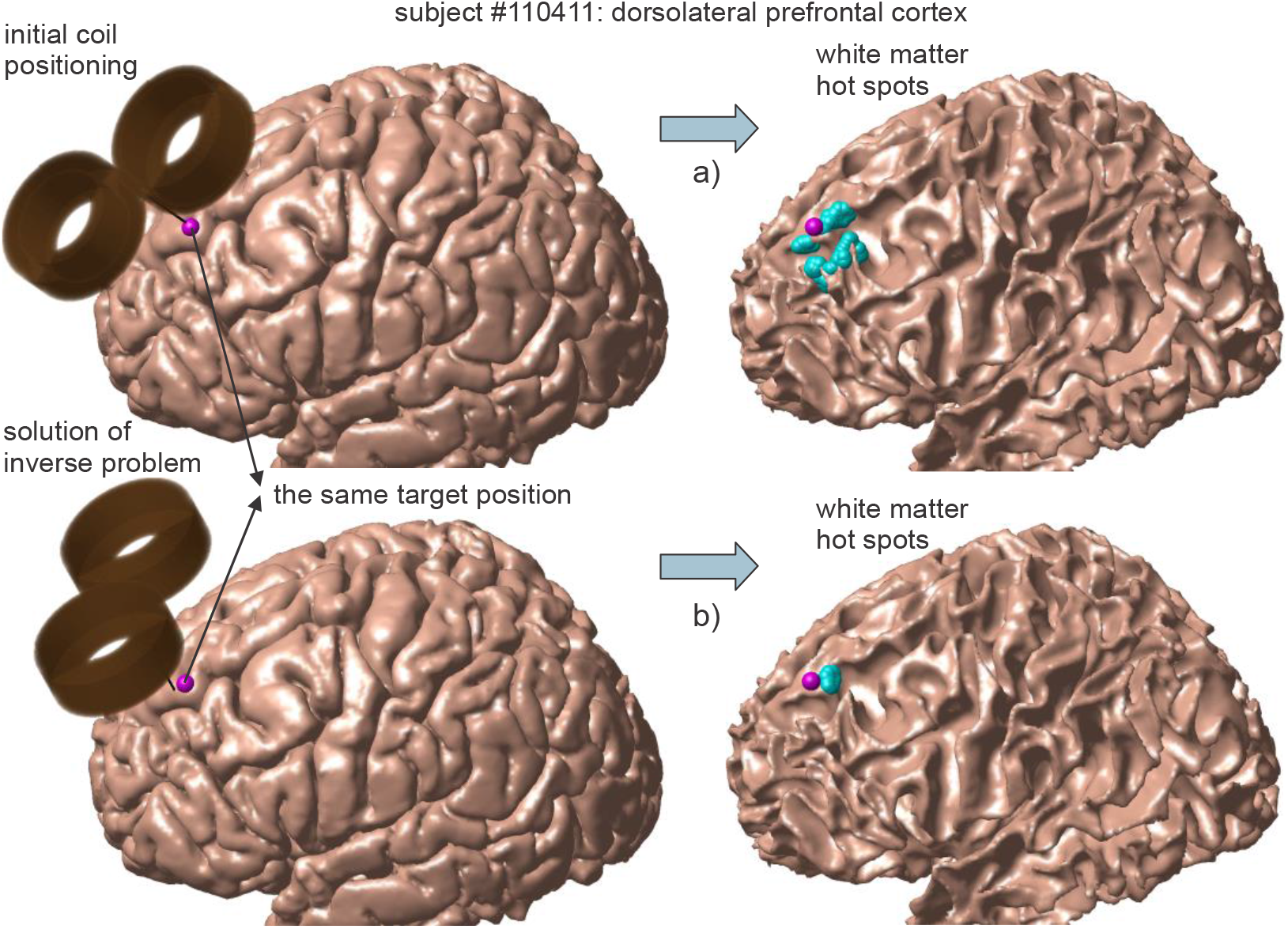
a) – Left: initial coil positioning 10 mm above the left dorsolateral prefrontal cortex of the left hemisphere for subject #110411 (GM surface is shown). The target position on the gray matter surface is marked by a magenta sphere. Right: white-matter “hot spots” – small blue balls drawn at the center of every WM facet where the absolute normal-field value is in the range of 80-100% of the maximum normal-field. b) – The same as a) but the coil position on the left is altered according to the inverse-problem solution.

### 3.3. Complete solution roster

Fig. 5 shows the complete inverse-problem solution data for sixteen subjects and for two targeted cortical areas. Fig. 5a indicates initial (darker color) versus final (lighter color) coil positions for each of the sixteen subjects while targeting the motor hand area; Fig. 5b gives the same result for the left prefrontal dorsolateral cortex.

**Fig. 5.**
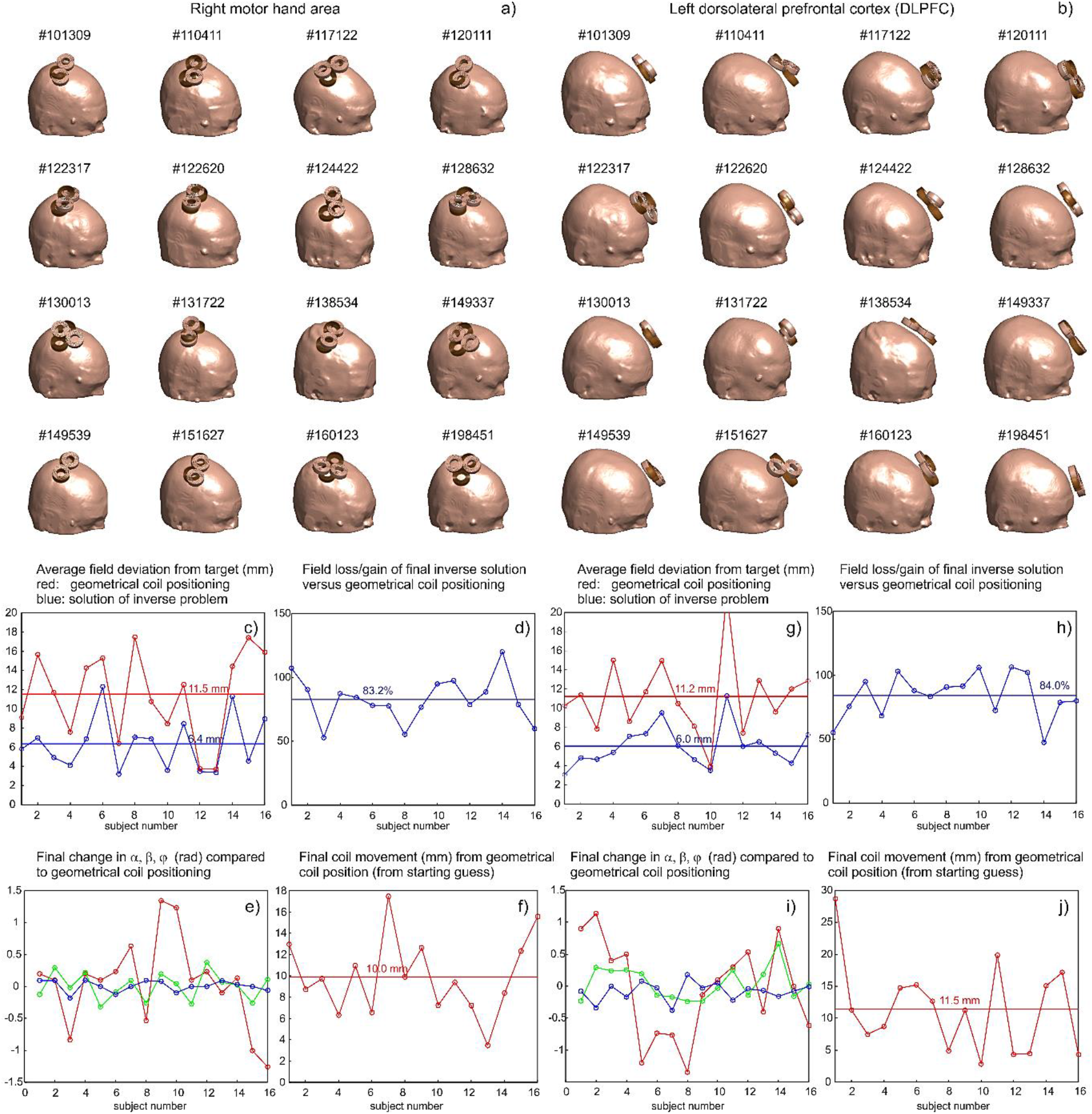
a) – Initial (darker color) versus final (lighter color) coil positions for each of the sixteen subjects while targeting the motor hand area. b) – The same result when targeting the left prefrontal dorsolateral cortex. c), g) – Cost function of Eq. (2) (initial-red versus final-blue) for two targeted cortical areas in mm. The average values of the cost function for the initial and final guess, respectively, are marked by two straight lines. d), h) – Field loss/gain percentage of the final inverse solution with respect to the maximum normal field at the inner cortical surface versus the initial guess. The average values of 83.2% and 84.0%, respectively, are indicated by two straight lines. e), i) – Changes in three rotation angles for the final solution as compared to the initial guess (red – *φ*, green – *α*, blue – *β*). f), j) – Deviation between the initial and final coil positions in mm; the average values of 10.0 mm and 11.5 mm, respectively, are indicated by straight lines.

Figs. 5c,g show the cost function of Eq. (2) (initial-red versus final-blue) for two targeted cortical areas, respectively. The units are mm. The average values of the cost function for the initial and final guess, respectively, are marked by two straight lines. They differ by approximately a factor of two in every case. Figs. 5d,h present field loss/gain percentage of the final inverse solution with respect to the maximum normal field at the inner cortical surface for the initial guess. The average values of 83.2% and 84.0%, respectively, are indicated by the straight lines. Figs. 5e,i indicate changes in three rotation angles for the final solution as compared to the initial guess (red – *φ*, green – *α*, blue – *β*). Figs. 5f,j report the deviation between the initial and final coil positions in mm; the average values of 10.0 mm and 11.5 mm, respectively, are indicated by straight lines.

### 3.4. Stability of the solution

The stability of the inverse problem solution is critical. It has been checked in every case using the following method. A cube in the ℜ^6^ search space centered around the final solution point was created, with three spatial sides of 3 mm each (as shown in Fig. 6a) and with three angular sides of 6 degrees, respectively. The cost function for all 729 points of the cube was then evaluated and averaged. After that, it was divided by the minimum cost function – the solution of the inverse problem – and the resulting relative average de-focalization was found as shown in Figs. 6b,c, respectively. Further averaging for all sixteen subjects was performed; it is shown in Figs. 6b,c by two straight lines, respectively. The final average values of 1.18 (118%) and 1.19 (119%) in Figs. 6b,c are remarkably close to each other and are both relatively close to one. This indicates an acceptable stability of the solution with regard to the position uncertainties observed for the navigated TMS (nTMS) [49],[50] and for robotic TMS where the corresponding values are approximately 3.5 mm (overall, 1.3 mm without coil–head spacing) and 3.5° [51] or 2 mm [52].

**Fig. 6.**
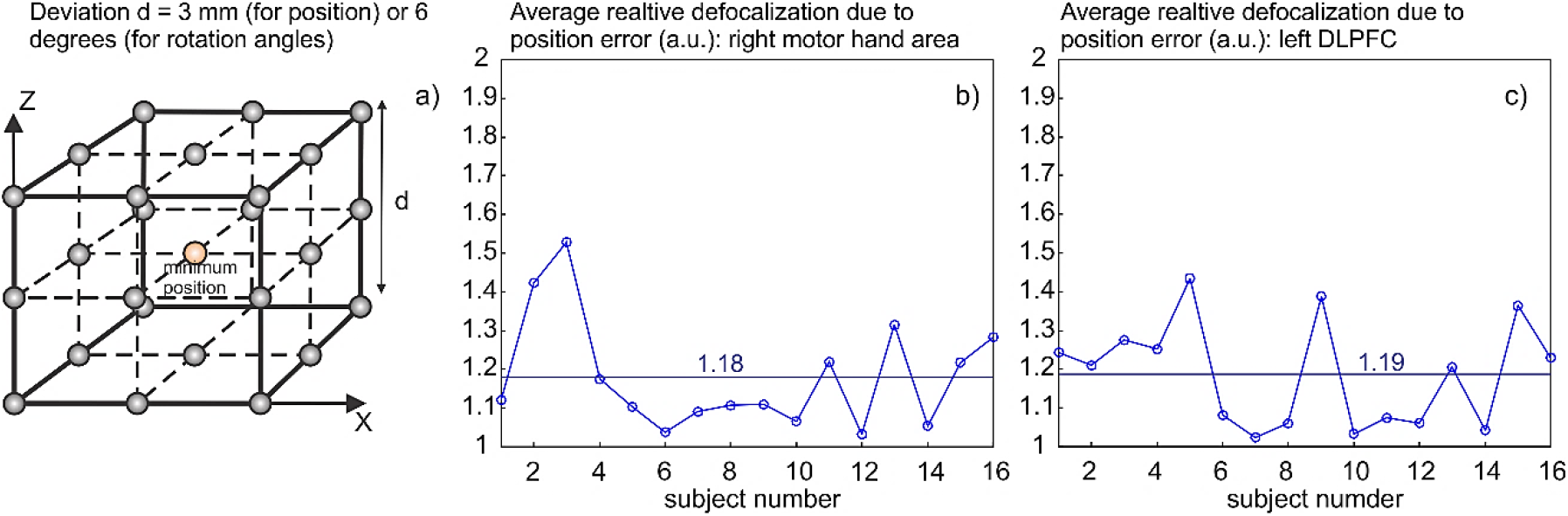
a) – A cube with 3 mm sides created for stability evaluation; a similar cube with sides of 6 degrees is created in the angular search space. b), c) – Relative average de-focalization versus the minimum cost function for the motor hand area and the prefrontal dorsolateral cortex, respectively. The result averaged for all sixteen subjects is shown by a straight line.

### 3.5 Scanning through the motor hand area. Potential for accurate mapping individual hand muscles

A separate experiment has been performed with the goal of establishing whether or not the method is capable of accurately scanning through the motor hand area. The results for subject #101309 of the Connectome database are given in Fig. 7. The approximate motor hand area of the right hemisphere is highlighted in pink. The magenta ball in Figs. 7a,b,c for the GM shell indicate the target position, which continuously moves along the precentral sulcus in the medial direction. Figs. 7d,e,f show the corresponding inverse problem solutions for the three cases considered. The spread of the intracortical normal electric field within 80-100% of the maximum normal field is indicated by small blue balls on the white matter surface. In all cases, this spread is minimal, suggesting that it may be possible to accurately differentiate motor representation of individual hand muscles. Volumetric fields of Fig.8 below exactly correspond to Fig. 6b,e, respectively. Prior relevant experimental studies were limited to two [53] and (not always entirely successfully) to three (abductor pollicis brevis, first dorsal interosseous, abductor digiti minimi) distinct hand muscles [54],[55]; one more recent relevant study is Ref. [18].

**Fig. 7.**
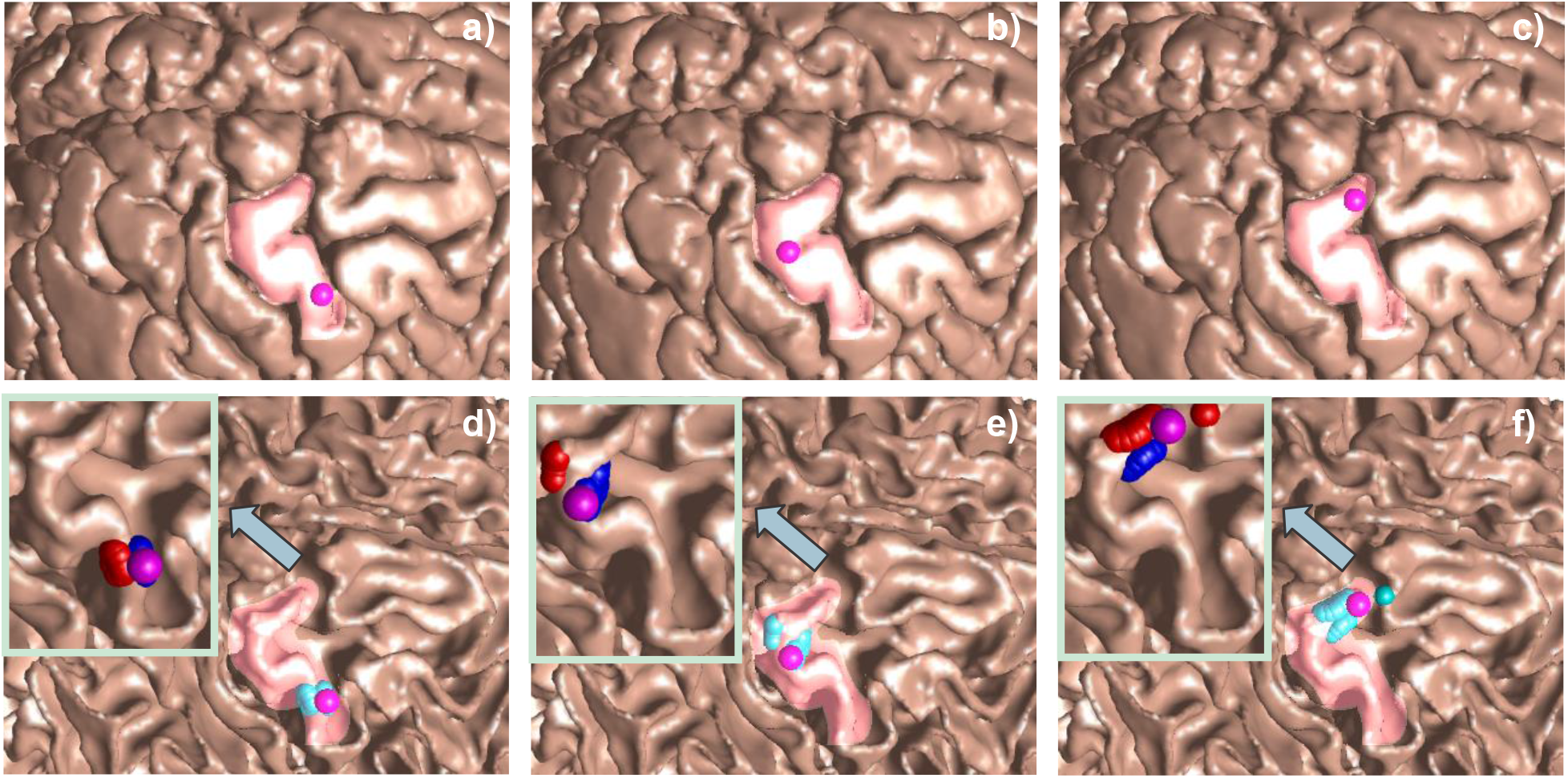
a-c) – Lateral-medial scanning through the motor hand area highlighted by a pink color for subject #101309 of the Connectome database with the moving target point marked by a magenta ball on the gray matter surface. d-f) – Computed inverse problem solution. Spread of the intracortical absolute normal electric field within 80-100% of the maximum absolute normal field is indicated by small blue balls on the white matter surface. The insets in d-f) show zoomed in *signed* field spread (positive-red or negative-blue normal field direction in Eq. (1)) corresponding to depolarization or hyperpolarization of the membrane potential, respectively. Volumetric fields of Fig.8 below exactly correspond to Fig. 6b,e.

## 4. Discussion

### 4.1. Stability correction

In four cases out of the thirty-two total cases reported in Fig. 5, the inverse problem solution has been corrected following a failed stability check. Those special cases can be divided into two groups. Subject #13 in Fig. 5c and subject #10 in Fig. 5g belong to the first group. Here, the initial guess is already very good: the average absolute deviation of the field maxima from the target point is below 4 mm. The inverse problem solution only marginally improved the focality but eventually became unstable. The relative average de-focalization depicted in Fig. 6 reached 1.8-2.0. Therefore, the iterative solution has been reverted to the first iteration that appeared to be stable in both cases, with the final stable result shown in Fig. 6b and Fig. 6c, respectively.

Two other cases (subjects 15 and 16 in Fig. 5c for the motor hand area) indicated a rather normal convergence, but the final solution also became unstable: the relative average defocalization depicted in Fig. 6 reached 1.7-1.9. In every case, the iterative solution has been reverted to a previous stable iteration (10^th^ iteration for subject 15 and 11^th^ iteration for subject 16 in Fig. 5c), with the final stable result shown in Fig. 6b.

### 4.2 Computational times

Although the exhaustive search algorithm employed to solve the inverse problem is safe, it requires large processing times. The total execution time including the stability check involves solving up to 10,000 forward problems per target position. Every such forward problem is also solved iteratively, using a generalized residual method or GMRES. The process is parallelized within MATLAB 2019a platform, using binary MEX FMM executables [31]. This results in approximately 12 hours of real time per one target position using an Intel Xeon E5-2698 v4 CPU (2.10 GHz) server, 256 GB RAM.

A better algorithm may start the iterative solution of every forward problem with the solution found for the neighbor position; this change may improve speed by a factor of 2-4. Additional speed improvement may require a Nelder-Mead simplex method [56] in ℜ^6^ or a variation of a generic algorithm [57], and better parallelization.

### 4.3 Extension to the total electric field. Possible correlation with the I wave activation sites

While only the normal component of the electric field at the inner cortical surface has immediately been concerned, we checked all final results with regard to the tangential and the total electric field distributions. It has been found that the inverse solution obtained for the normal field may quite well correspond to the global maximum of the total electric field and to the maximum of the field gradient within the entire gray matter volume, which are associated with I wave excitation.

To demonstrate this, Figs. 8a,b,c left present transverse, coronal, and sagittal planes passing through the point of the maximum normal field at the white-gray matter interface and superimposed onto the corresponding NIfTI slices for the targeted hand knob area of the right precentral gyrus of subject #101309. This figure exactly corresponds to the inverse-problem solution for the targeting result of Fig. 7b,e; only the field plots are drawn differently.

**Fig. 8.**
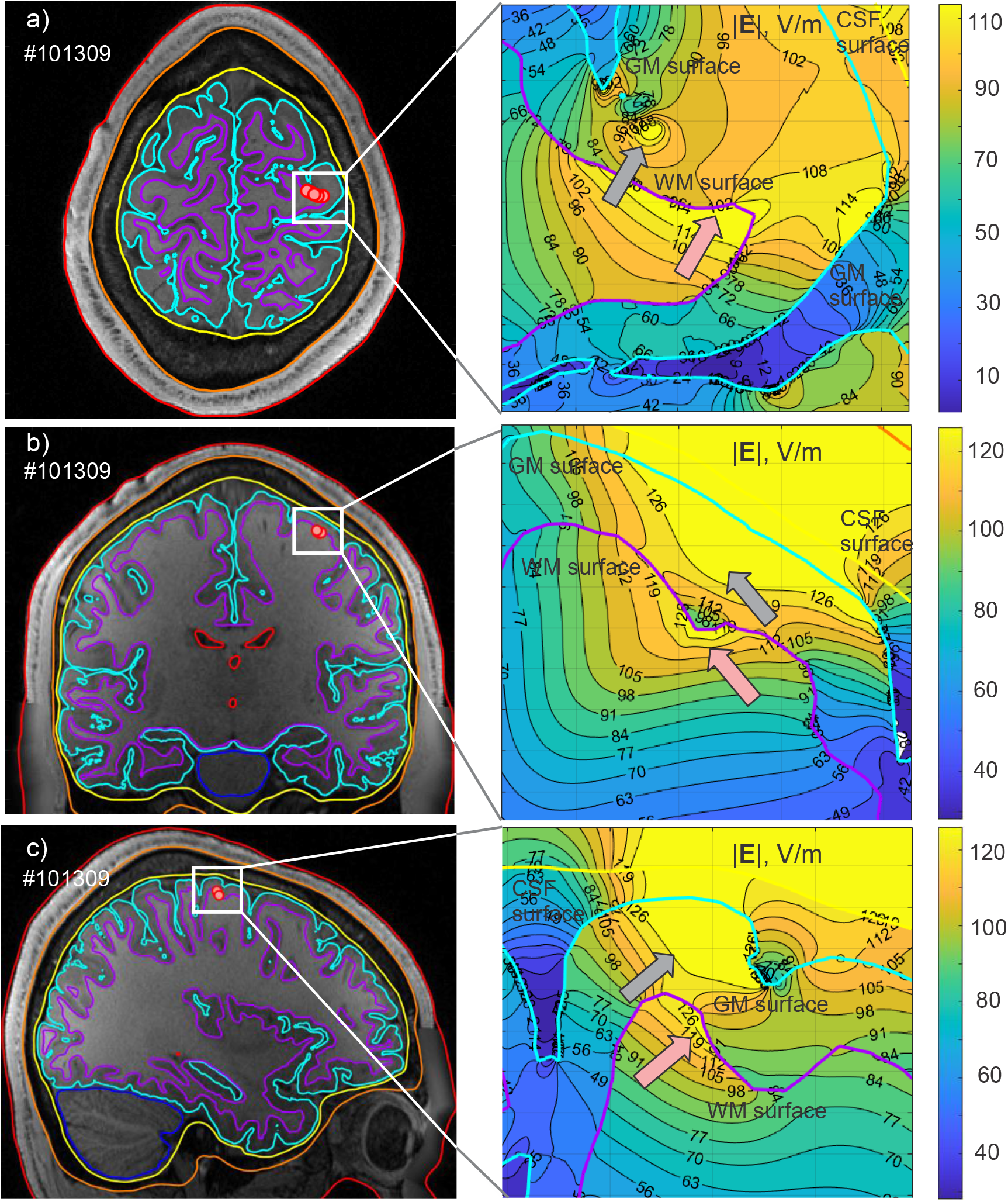
a,b,c) – Left: transverse, coronal, and sagittal planes passing through the point of the maximum normal field at the white-gray matter interface and superimposed onto the corresponding NIfTI slices of subject #101309 for the targeted hand knob area of the right precentral gyrus. This figure exactly corresponds to the inverse-problem solution for the targeting result of Fig. 7e; only the field plots are drawn differently. The red dots show the centers of intersected white matter facets where the absolute normal-field value is in the range of 80-100% of the maximum normal-field value. a,b,c) – Right: volumetric total electric field (field magnitude) distributions in V/m within the small white rectangles in Figs. 8a,b,c left. Maximum of the field discontinuity at the inner cortical surface is approximately indicated by pink arrows; maximum of the total TMS field (and apparently its gradient) observed in the gray matter volume is approximately indicated by grey arrows.

The red dots in Figs. 8a,b,c show the centers of intersected white-matter facets where the absolute normal-field value is in the range of 80-100% of the maximum normal-field value. At the same time, Figs. 8a,b,c right show volumetric total electric field (field magnitude) distribution within the small white rectangles in Figs. 8a,b,c left. One observes that the maximum total field magnitude (indicated by a grey arrow) and the associated maximum field gradient occur near the normal-field maximum (marked by a pink arrow). Similar results have been observed in other cases.

## 5. Conclusions

The inverse TMS problem considered in this study is the limited in scope yet computationally successful attempt to produce more focal TMS fields. We restrict ourselves by the “hot spot” (or spots) of the normal electric field at the inner cortical surface (the white-gray matter interface) and its immediate vicinity. These hot spots, biophysically corresponding to the earliest D-wave activation, must be located near a known target location. This is done by altering coil position and rotation via the inverse-problem solution. The search space is a six-dimensional space of three coil coordinates and three rotation angles. The numerical solution is determined by the unique gyral pattern of a subject and requires a detailed individual cortical surface segmentation. The practical realization of the solution relies on accurate coil positioning via robotic or navigated TMS.

For 16 subjects and 32 distinct target points (16 for the motor hand area and 16 for the prefrontal dorsolateral cortex), the inverse-problem solution decreases the mean deviation of the maximum field (80-100%) domain from the target point by approximately a factor of 2 on average, as compared to the initial geometrical coil positioning. This results in final deviation values that approach 6 mm on average. The corresponding reduction in the maximum-field area (the squared measure of deviation) is expected to quadruple.

The rotation angle about the coil axis is the most significantly altered parameter. The coil movement from the initial position is approximately 10 mm on average.

Stability with respect to perturbations in coil position and angle(s) is checked and corrected (if applicable) for every solution. The relative average de-focalization is less than 1.2 when the coil position accuracy is better than or equal to 3 mm and coil rotation accuracy is better than or equal to 6 degrees. The inverse-problem solution also causes electric-field strength loss as compared to the initial geometrical coil positioning. This loss is 16% on average.

The solution obtained may correspond to the nearby global maximum of the total electric field and to a maximum of the field gradient within the entire gray matter volume, which are associated with later I wave activation at low suprathreshold TMS intensities or elsewhere.

